# Three-dimensional histology reveals dissociable human hippocampal long axis gradients of Alzheimer’s pathology

**DOI:** 10.1101/2023.12.05.570038

**Authors:** Diana Ortega-Cruz, Kimberly S. Bress, Harshvardhan Gazula, Alberto Rabano, Juan Eugenio Iglesias, Bryan A. Strange

## Abstract

**INTRODUCTION:** Three-dimensional (3D) histology analyses are essential to overcome sampling variability and understand pathological differences beyond the dissection axis. We present Path2MR, the first pipeline allowing 3D reconstruction of sparse human histology without an MRI reference. We implemented Path2MR with post-mortem hippocampal sections to explore pathology gradients in Alzheimer’s Disease.

**METHODS:** Blockface photographs of brain hemisphere slices are used for 3D reconstruction, from which an MRI-like image is generated using machine learning. Histology sections are aligned to the reconstructed hemisphere and subsequently to an atlas in standard space.

**RESULTS:** Path2MR successfully registered histological sections to their anatomical position along the hippocampal longitudinal axis. Combined with histopathology quantification, we found an expected peak of tau pathology at the anterior end of the hippocampus, while amyloid-β displayed a quadratic anterior-posterior distribution.

**CONCLUSION:** Path2MR, which enables 3D histology using any brain bank dataset, revealed significant differences along the hippocampus between tau and amyloid-β.

## 1. Background

A wealth of information regarding region-specific cellular and pathological features has been obtained from human post-mortem brain specimens^1–4^. For this purpose, brains are typically cut into slices, from which smaller, thin sections are sampled and analyzed under the microscope. The ensuing observations are assigned to the anatomical structure of origin and compared across subjects. However, since in standard practice sampled regions are visually identified and manually processed, histological sections from different subjects rarely originate from the same exact brain position. This results in a lack of anatomical generalizability of the sophisticated cellular and pathological quantifications which are attracting increasing interest^5–7^. Identifying the precise structural location of each section would allow performing anatomically relevant comparisons of histological observations. Moreover, this would enable pooling data from different individuals into a common space to study histological measurements beyond the sectioning plane.

To achieve such fine-grained mapping, three-dimensional (3D) histology reconstruction approaches have been developed, which rely on image registration to align consecutive histological sections^8^. Without a reference of the original shape, inferring the 3D volume conformation represents a challenge, and often leads to artifacts such as z-shift (accumulation of errors along the stack) and the “banana effect” (straightening of curved structures)^8^. Several strategies are available to optimize reconstruction outcome^8–12^. These methods have been applied for brain reconstruction using animal^13–15^ and human histology^9,16^. However, they require dense, serial sectioning of the whole brain or structure of interest to ensure accurate representation of its 3D configuration^17^. Such dense histological sampling is rarely performed in routine brain bank procedures and may be unfeasible due to time limitations or sample requests for other purposes.

Alternative efforts towards histology reconstruction have relied on Magnetic Resonance Imaging (MRI) as a 3D reference. Using subject-specific *in vivo*^18,19^ or *ex vivo*^20–24^ scans, these approaches overcome shape uncertainty by registering sections to the MRI volume. With this strategy, serial histology reconstruction allows accurate unbiased registrations to an atlas. Additionally, other MRI-based methods have enabled reconstruction of sparse histological images^18,19,25–27^. Although entailing less accurate slice-to-volume registrations^28^, this option avoids dense sampling of the whole specimen. Unfortunately, *in vivo* MRI is only available for special cases or in planned follow-up initiatives such as the Alzheimer’s Disease Neuroimaging Initiative (ADNI)^29^. Similarly, many brain banks have no access to *ex vivo* scanning due to financial and logistic constraints. As these approaches rely on highly specialized setups and equipment, their applicability to the large histological datasets generated in brain banks and clinical facilities is limited.

Here, we present Path2MR, a histology mapping pipeline that enables 3D studies using sparse sampling and without a specific MRI reference. Our strategy uses blockface images of brain hemisphere slices to recover the 3D structure by geometrical stacking. Then, deep learning is employed to obtain a 1mm^3^ resolution 3D prediction of the hemisphere with MRI-like contrast, which is subsequently registered to the widely used Montreal Neurological Institute (MNI) atlas. Histological sections are then registered to the corresponding coronal slice within the reconstructed hemisphere, and deformations are concatenated to register them to MNI standard space.

We demonstrate the applicability of Path2MR by analyzing pathological gradients of Alzheimer’s Disease (AD), for which neuropathology serves as ground truth for diagnosis and biomarker validation^30^. Specifically, we explore the distribution of tau and amyloid-β (Aβ) pathologies along the anterior-posterior axis of the hippocampus in 26 patients with no co-pathologies aside from AD. We take advantage of the spatial variability inherent to histological sampling to achieve a fine depiction of this axis using only three sections per subject. Our results show an anterior-posterior gradient of hippocampal tau pathology, consistent with previous work using *ex vivo* MRI and dense sampling^31^. In contrast, Aβ pathology displays a quadratic-shaped distribution, with more variable patterns of deposition across hippocampal subfields. Our 3D histology pipeline is widely applicable to any prospective or retrospective brain bank histology dataset.

## 2. Methods

Path2MR is summarized in Figure 1, and steps are detailed in Sections 2.1 to 2.5. Steps described in Sections 2.1-2.4 (3D reconstruction and registration to MNI) are performed once for every subject, while histology registration (Section 2.5) is performed independently for each section included in population analyses. This pipeline is agnostic to the brain processing procedures. As many brain banks limit histology to one hemisphere, the pipeline has been developed for single hemisphere reconstruction; however, it is easily adaptable to 3D reconstruction of the whole brain. Steps described in Sections 2.1 and 2.2 are available as part of the neuroimaging suite FreeSurfer, and steps from Sections 2.3-2.5 are publicly available at https://github.com/ortegacruzd/Path2MR.

**Figure 1.**
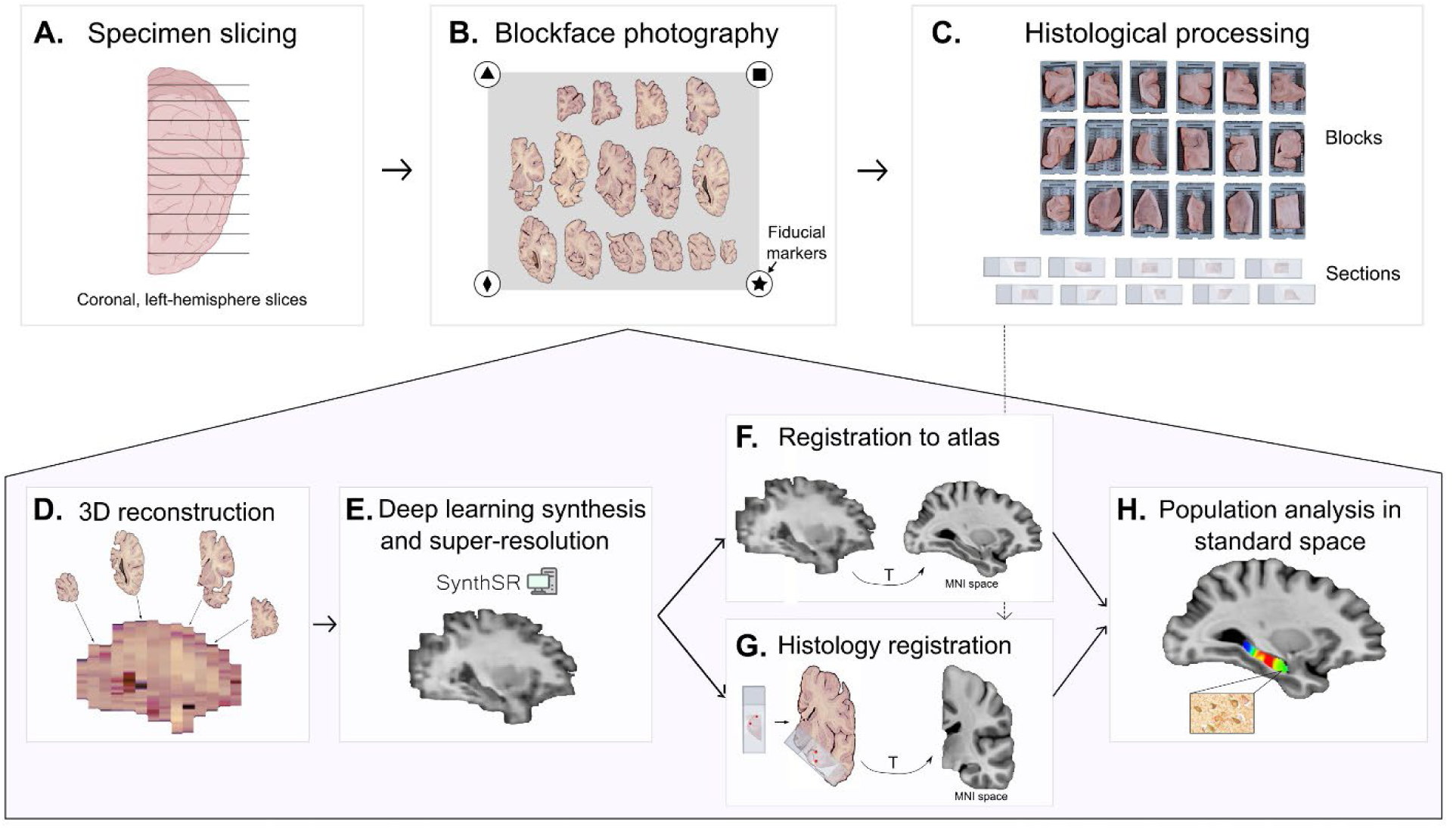
Summary of Path2MR. Steps in the top row include tissue processing and photography (Section 2.1), and the bottom shaded pentagon includes steps of the reconstruction method (Sections 2.2 to 2.5). Briefly, the fixed specimen is: **A.** sliced (coronally and unilaterally in our case), **B.** photographed including fiducial markers, and **C.** dissected for histological processing. The 3D reconstruction method uses blockface photographs as the starting point (**D.**, Section 2.2), followed by deep learning synthesis of a 1mm^3^ resolution prediction of the hemisphere (**E.**, Section 2.3). To enable population analyses, an atlas in MNI space is registered to this hemisphere prediction (**F.**, Section 2.4). Linear and non-linear transforms obtained at this step (represented with a T) are used after registration of histology sections to the 3D reconstruction, for which manual initialization is used (**G.**, Section 2.5), to in turn register histology to MNI. Subsequently, any histology-derived measure can be compared for population analysis in standard space (**H.**). 3D: three-dimensional. MNI: Montreal Neurological Institute.

### 2.1 Blockface photography

The procedure for blockface photography of specimen slices has been recently described^32^ and photograph processing is available in FreeSurfer 7.4^33^ (and subsequent versions). As shown in Fig. 1A, the specimen destined for histology should be cut by an experienced pathologist into slices of regular thickness (ideally under 10 mm to optimize the reconstruction result). Specimen slices are placed on a flat (preferably dark) surface including four fiducial markers on the corners of a rectangle of known dimensions (Fig. 1B). All slices are placed in the same orientation (either anterior or posterior), and photographs are then taken with homogeneous lighting. The fiducials are automatically detected with Scale Invariant Feature Transform (SIFT)^34^ and Random Sample Consensus (RANSAC)^35^, or alternatively, four reference points of known distance between them can be manually selected within each photograph. The distance between fiducials or reference points is used to compute a perspective transform that corrects geometric distortion of the photographs, while calibrating their pixel size^32^. This correction overcomes the variable angles of different elements in the image to the camera, and can also adjust for potential variations in camera distance across photographs. After photography, blocks are sampled from the slices for histological processing (Fig. 1C).

### 2.2 3D photograph reconstruction

From corrected photographs, slices are segmented from the background using automatic color thresholding (easier with dark backgrounds) or manual correction (if background is light-colored) in any image editing software, such as the open-source package GIMP (https://www.gimp.org). Then, we define the order of the slices using a simple graphic user interface (GUI), also available in FreeSurfer 7.4.

Next, the slices are reconstructed into a 3D volume^32^ (Fig. 1D) using a joint registration framework for MRI-free reconstruction^36^. This framework takes advantage of prior knowledge about brain slice thickness and uses an MRI atlas (MNI) as 3D reference volume. The orientation of slices in the image, either anterior (default) or posterior can be specified, resulting in interchangeable results.

### 2.3 Deep learning synthesis

The reconstructed volume is aligned to the reference MNI atlas only linearly, so accurate mapping requires subsequent nonlinear registration. In order to increase the accuracy of nonlinear alignment, it is desirable to synthesize a 1 mm isotropic synthetic MRI (Fig. 1E) from the 3D reconstructed volume^37^. For this purpose, we use SynthSR^38,39^, which we finetuned to adapt the method to the features of slice photography: absence of cerebellum or brainstem; single hemisphere; and different brightness variations for every coronal slice. As shown previously^38^, SynthSR confers reliable results across brain structures, and was trained using data from subjects with varying degrees of atrophy, making it robust to severely atrophied brain specimens.

### 2.4 Registration to MNI

As shown in Figure 1, SynthSR produces a 1 mm isotropic volume with T1-weighted MRI-like contrast. We register the T1-weighted version of the MNI atlas to this volume (Fig. 1F) using NiftyReg^40^, an open-source software for linear and non-linear registration of medical images (http://cmictig.cs.ucl.ac.uk/wiki/index.php/NiftyReg). NiftyReg is a widely used tool showing highly accurate results, with a validated performance compared to other registration methods^41,42^. Registration includes a linear step (function *reg_aladin*) followed by a non-linear diffeomorphic registration, using 15mm control point spacing and a local normalized cross correlation objective function with 4mm gaussian window (function *reg_f3d -vel -sx -15 --lncc 4.0*).

### 2.5 Histology registration and pathology map computation

Direct slice-to-volume registration of a histological section to MNI space is extremely ill-posed and ambiguous, particularly for nonlinear registration^28^. To circumvent this challenge, our pipeline takes advantage of prior knowledge of the brain slices from which tissue blocks (and histology sections) were derived. This correspondence is used to register each histological section to its approximate location and rotation in the brain slice. Manual initialization is performed by selecting two anatomical gyrification landmarks (e.g., for hippocampal sections, the hippocampal fissure and the border of the temporal horn) in both the histology section and its corresponding hemisphere slice (Fig. 1G). To account for block trimming during sectioning, if slice photos and tissue blocks were obtained from the anterior side, we shift the initial coronal position of the section 2.5mm posteriorly. This shift was chosen assuming histology sections were obtained from the center of the tissue block (commonly 5mm thick), entailing an error of ±2.5mm in registration initialization. Finally, the MNI atlas registered to subject-specific space (Section 2.4) is used to enhance registration accuracy in the coronal plane based on gradient magnitude correlation, within a range of ±10mm. This last step is particularly useful if working with thicker dissection blocks, which would entail a higher initialization error.

To register histology to atlas space, the coronal level of each section resulting from the prior registration step is used as a reference. To model uncertainty in the anterior-posterior direction as the main source of error in this pipeline, we use kernel regression, whereby kernels account for both the density of the data (for interpolation purposes) and registration error. To that end, a gaussian distribution around each section’s resulting coronal position is obtained, serving as a position probability function. Linear and nonlinear deformations obtained in step 2.4 are then concatenated to nonlinearly deform gaussian distributions to MNI space (Figure 2). Subsequently, a weighted average of measured pathology burdens (or any other histological measurement) is obtained, using their gaussian position distributions as weights. A mask of the brain structure of interest is then applied to the resulting average burden maps, thereby enabling population analyses (Fig. 1H).

**Figure 2.**
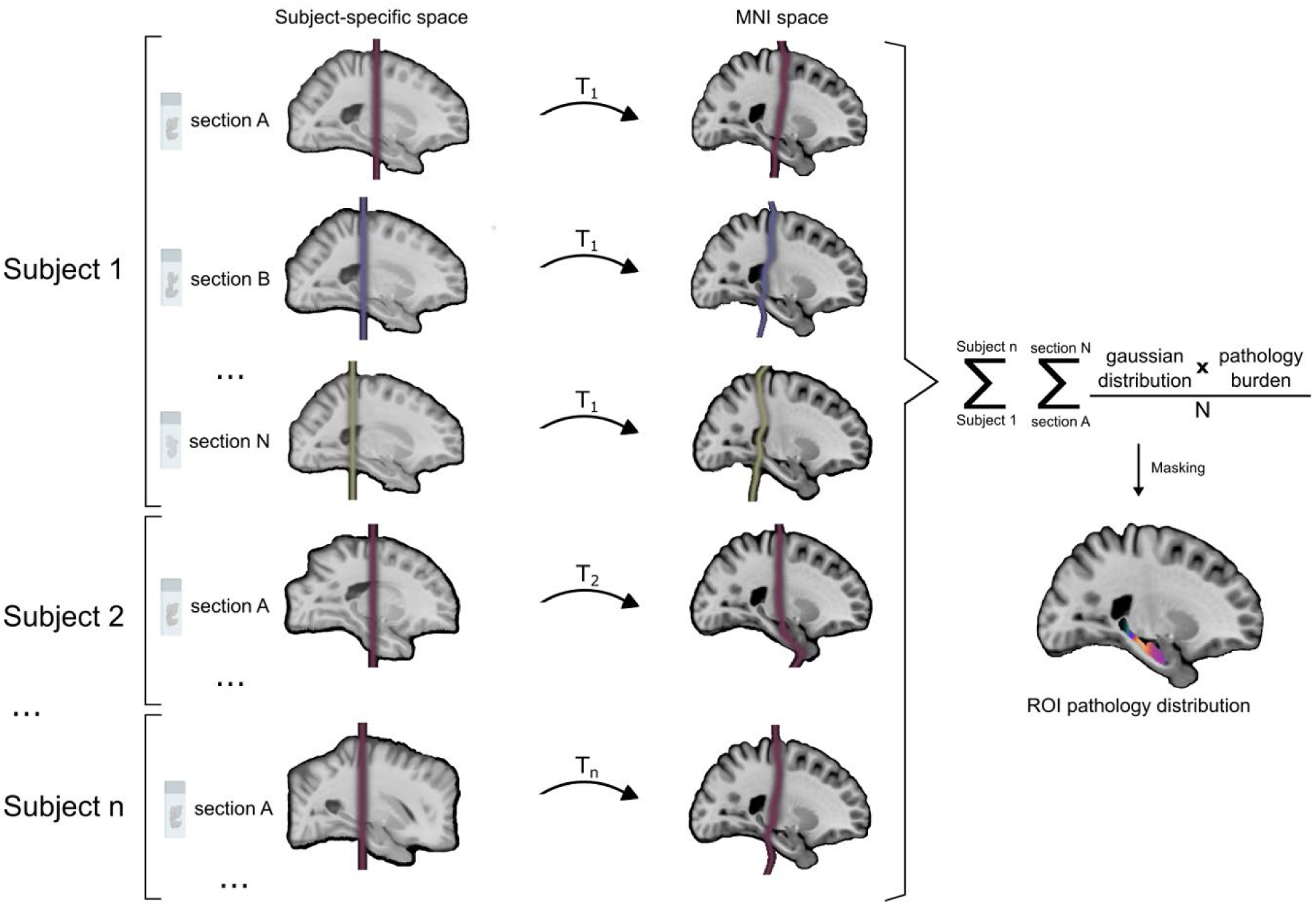
Steps followed to perform population analysis of histology in standard space. First, gaussian distributions are obtained around the position of each section in subject-specific space. These are then registered to MNI space using transforms derived from registration of the atlas to the synthetic MRI for each subject (T1, T2,…Tn). The gaussian position distribution of each section in MNI space is multiplied by its quantified pathology burden and normalized by the total number of sections included for that subject. Finally, burden maps for all sections and subjects are added and masked to obtain the pathology distribution of the population within the region of interest (ROI). MNI: Montreal Neurological Institute; n: number of subjects; N: number of sections per subject; ROI: region of interest.

To obtain MNI coronal coordinates for Path2MR validation compared to visual positions, after registration to MNI, gaussian distributions of each histology section were applied a mask of the hippocampus. Within each masked distribution, the coronal slice with highest number of high-intensity voxels (equal to 1) was selected as the coronal coordinate of the section (the center of the distribution). Intensity measures per coronal plane were derived from masked distributions and pathology maps using commands from FSL software^43^, specifically *fslslice*, *fslmaths* and *fslstats*. Throughout the manuscript, the uncus was used as reference for the transition from hippocampal head to body, delimiting y=-22 as the first MNI coronal coordinate within the hippocampal body^44^.

### 2.6 Experimental donor cohort

To evaluate the applicability of the presented pipeline, we used samples from donors with dementia from the BT-CIEN brain bank (Madrid, Spain). Standard diagnostic procedures at BT-CIEN^45^ include qualitative assessment of global and medial temporal lobe (MTL) atrophy (0-3) immediately after extraction. Subsequently, the left hemisphere is fixed in 4% phosphate-buffered formaldehyde and cut into 10mm-thick coronal slices for tissue block dissection, while the right hemisphere is frozen. Neuropathological evaluation is performed according to published criteria for AD^46^, vascular pathology^47^, presence of Lewy Bodies^4^, LATE^48^ and hippocampal sclerosis of aging (HS)^49^.

To minimize heterogeneity from co-pathology, we included all patients with a neuropathological diagnosis of “pure” non-familial AD according to the following criteria: i) high AD neuropathologic change (ADNC)^46^; ii) no Lewy Bodies, LATE nor classical HS, and low or no vascular pathology; iii) extent of global and MTL atrophy lower than the maximum (i.e., score 0-2), as neuropathologic criteria could not be reliably evaluated in samples from extremely atrophic cases (score of 3). Intermediate ADNC cases with no co-pathology commonly present milder clinical profiles^50^ and are therefore rare in dementia autopsy series. Out of all donations received at BT-CIEN between 2007 and 2020, a total of 26 amnestic dementia patients complied with these criteria and were included in the study.

Reconstruction was performed from retrospective photographs of stored hemisphere slices previously processed for histology. For all subjects, reconstruction was carried out using anterior blockface photographs. If any slice had missing information due to tissue dissection, with the posterior side of its consecutive slice being intact, the latter was replaced by the former (horizontally flipped) prior to segmentation from the background (Section 2.2) using GIMP 2.10.28.

### 2.7 Histology and quantification

Given the extensively studied vulnerability of the hippocampus to AD pathology, we employed Path2MR to analyze the distribution of tau and Aβ pathologies along the hippocampal longitudinal axis. As shown in Figure 3A, we used histology sections obtained from three coronal blocks of the left hippocampus: hippocampal head, body and tail. Blocks were processed using the HistoCore PEARL (Leica Biosystems), placed within paraffin, and the Microm HM355S microtome (ThermoFisher) was used to obtain 4μm-thick sections. Immunohistochemistry was carried out using primary antibodies for tau AT100 (ThermoFisher Invitrogen, code: MN1060, 1:100 dilution) or Aβ (Dako, code: M087201-2; 1:40 dilution), together with hematoxylin. For each pathology, analyses were designed to include a total of 78 sections (three per subject). Out of these, nine blocks could not be included in tau analysis due to dim staining or tissue unavailability, and six could not be obtained for Aβ staining, resulting in 69 and 72 sections included, respectively. For histology registration (Section 2.5), a 1X digital image of each section was obtained using a Nikon Coolscan V-Ed scanner. The anatomical position of each section was determined by KSB and AR, blind to Path2MR outcome. This was performed based on the micro-atlas by Mai *et al.*^51^, determining the distance to the anterior commissure (AC) of each section (ranging from 10.7 to 39.5 mm) by visual inspection under the Nikon 90i light microscope.

**Figure 3.**
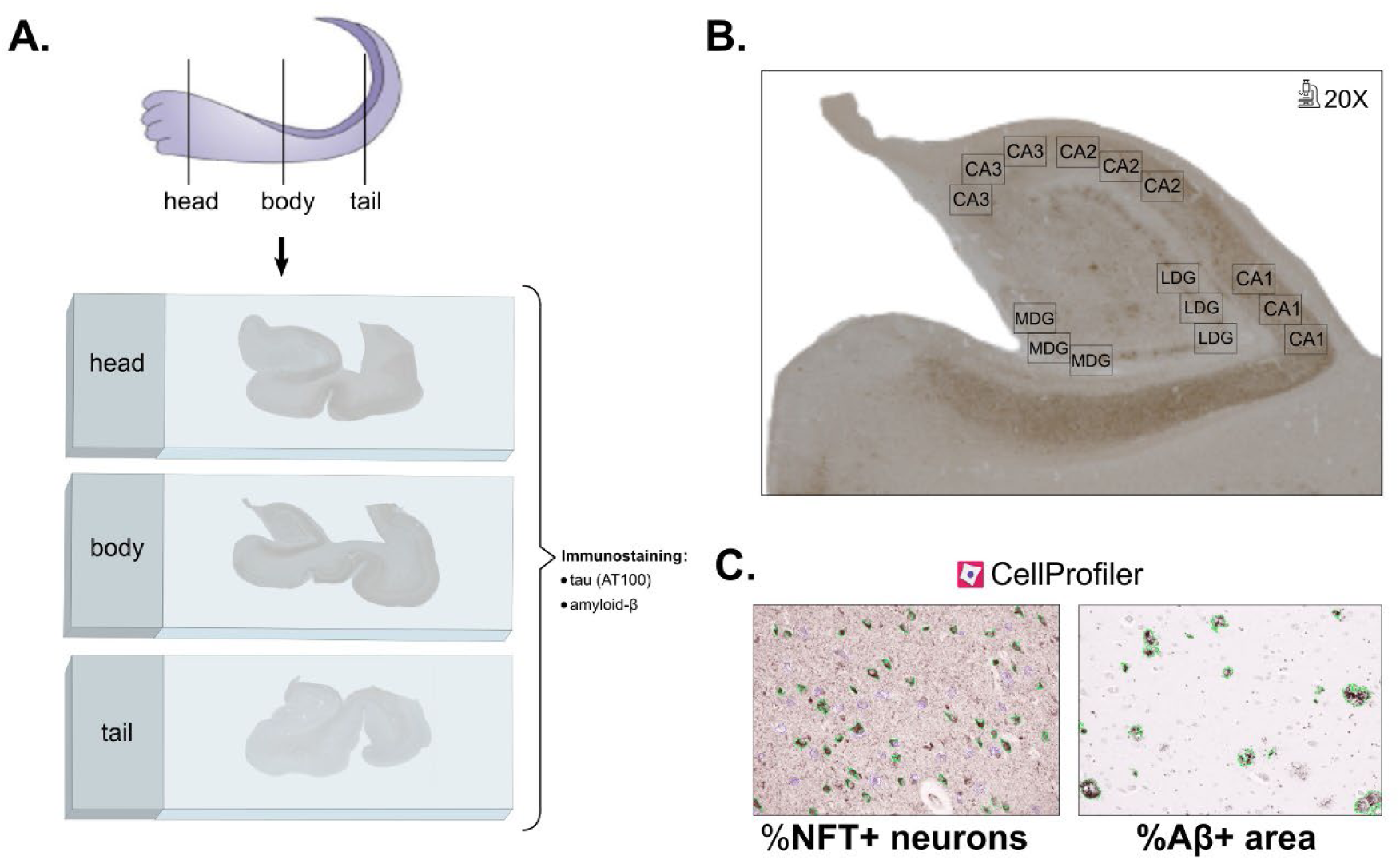
Summary of steps carried out for AD pathology quantification along the hippocampal longitudinal axis. **A.** Three anatomical blocks were sectioned for histology: head, body, and tail of the hippocampus. Sections from each of these levels were stained with antibodies against tau (AT100) and Aβ. **B.** Three micrographs at 20X magnification were taken per subfield: CA1, CA2, CA3, medial (MDG) and lateral dentate gyrus (LDG). **C.** Each 20X micrograph from sections stained against tau or Aβ was quantified with CellProfiler 4.0.7, measuring the percentage of NFT+ neurons (outlined in green, versus NFT-neurons outlined in purple) or area covered by Aβ plaques (outlined in green), respectively. Aβ: amyloid-β; LDG: lateral dentate gyrus; MDG: medial dentate gyrus; NFT: neurofibrillary tangles.

To obtain an average measure of tau neurofibrillary tangle (NFT) and Aβ burden per section, we measured these in five hippocampal subfields: CA1, CA2, CA3, as well as the medial (MDG) and lateral (LDG) regions of the dentate gyrus (Figure 3B). The selected CA1 region included the medial portion of the subfield (with proximal meaning close to CA2, and distal close to the subiculum). Three micrographs were acquired in each subfield using a Nikon 90i microscope with 20X magnification. Pathology burden was quantified using CellProfiler 4.0.7, performing segmentation based on the color of the stain followed by intensity correction to homogenize across micrographs. Then, in tau images, the following items of interest were identified based on size: NFT+ neurons (mature tangles, covering the whole neuronal soma) and hematoxylin-stained nuclei of NFT-neurons. In Aβ images, diffuse, cored, cotton-wool or coarse-grained plaques were identified, without separate classification of each type. As previously described, tau burden was measured as the percentage of NFT+/total neurons in the image^52^, and amyloid burden was measured as the area percentage covered by Aβ plaques^53^ (Figure 3C). As neuron size differs between pyramidal CA layers and granular DG layers, independent projects were used for tau quantification in CA and DG subfields. To normalize counts by layer area, as CA images only included the pyramidal layer, the whole area of the image was used. Conversely, DG images included the neuron-containing granular layer and neighboring (polymorphic and molecular) layers, and thus segmented neurons were dilated (to a size of 15 pixels) and merged to delimit the area of the granular layer. Our CellProfiler projects for tau and Aβ quantification are also available at https://github.com/ortegacruzd/Path2MR. Subfield burden was obtained as the average between the three micrographs within a subfield, and global section burden was obtained as the average among the five subfields.

### 2.8 Ground truth experiment

To validate Path2MR reconstruction performance, we used MRI scans from the Human Connectome Project (HCP)^54^ as ground truth dataset, adapting methodology by Gazula *et al.*^32^. T1 and T2 sequences (slice thickness of 0.7mm) from 100 subjects were first skull stripped using FreeSurfer, followed by extraction of the left hemisphere. The T1 was used as ground truth structure, and the T2 was used for reconstruction (Figure S1). To that end, we simulated 10mm-thick dissection photographs from T2 hemispheres by extracting one coronal slice every 14 (0.7mm*14=9.8mm spacing). To mimic variability in experimental data, each extracted slice was applied a distortion transform T_dist_ that comprised random rotations (within ±20°), translations (within ±0.5 pixels in both axes), shearings (within 10% in both axes), and smooth illumination fields. Larger random translations were not required, since 3D reconstruction is initialized by matching centers of gravity of the slices. Distorted T2 slices were then used for 3D reconstruction, deep learning synthesis and nonlinear MNI registration (Path2MR steps in Sections 2.2, 2.3 and 2.4, respectively). Finally, the T1 hemisphere was also applied a 3D random rigid transform (rotation within ±30° and translation within ±20mm) and nonlinearly registered to MNI using the same configuration as in Section 2.4 .

On one side, MNI registration of the T1 ground truth hemisphere yielded a deformation transform D_1_. On the other side, reconstruction of T2-derived slices yielded a restoration transform T_rest_, and after SynthSR processing, MNI registration of the resulting isotropic scan yielded a transform D_2_. We can then compare D_1_ with the composition: T_dist_ ∘ T_rest_ ∘ D_2_. In an ideal scenario, T_dist_=T_rest_^-1^ such that the two cancel each other, with SynthSR generating perfect isotropic images and D_1_ and D_2_ being identical. In practice, mistakes are made in the 3D reconstruction (which affects the error directly) and in deep learning synthesis (which affects the error indirectly, via impact on the NiftyReg registration). Therefore, the error was computed as the module of the difference between D_1_ and D_2_, averaged across non-zero voxels in every reconstructed slice.

### 2.9 Statistics

Statistical comparisons and associated plots were performed using RStudio 1.4.1106, including the following packages: *ggpubr*, *tidyverse*, *readxl*, *rstatix* and *viridis*. Tau and Aβ distributions along the longitudinal hippocampal axis were compared using Kolmogorov-Smirnov test. To find the most appropriate fit to the data, we used linear, quadratic, and cubic functions and selected the fit with highest adjusted R^2^ and lowest Akaike information criterion (AIC) value. Pathology burdens were compared between subfields using repeated measures analysis of variance (ANOVA) and post-hoc pairwise T tests.

### 2.10 Ethical approval

The BT-CIEN brain bank is officially registered by the Carlos III Research Institute (Ref: 741), by which donation is carried out under informed consent by a relative or proxy. BT-CIEN procedures, have been approved by local health authorities of the Madrid Autonomous Community (Ref: MCB/RMSFC, Expte: 12672). The study of these data was independently approved by the Ethics Committee of the Universidad Politécnica de Madrid (N° Expte: 2021-062).

## 3 Results

### 3.1 Validation of reconstruction performance

First, the accuracy of Path2MR in recovering the original structure of the specimen was quantitatively assessed using scans from 100 subjects from the Human Connectome Project. T1-weighted scans were used as ground truth, and corresponding T2-weighted scans were used to simulate 10mm-thick slices for 3D reconstruction, deep learning synthesis and registration to MNI (Figure S1). Resulting deformations were compared at every voxel with those obtained from MNI registration of the T1 ground truth, revealing an average 3D error of 3.27±0.42 mm. Evaluating separate contributions from each axis, we found an error of 1.22±0.18 in medial-lateral, 1.71±0.33 in superior-inferior, and 1.91±0.26mm in the anterior-posterior direction. The comparable error between the three axes, while dealing with discrete sampling in the anterior-posterior axis, shows that Path2MR steps prior to histology registration achieve an accurate estimation of the specimen’s 3D structure.

### 3.2 Validation of histology localization

Together with reconstruction performance, the reliability of results obtained with Path2MR depends on its ability to accurately localize histologic sections. To validate this step using our hippocampal histology dataset, we compared Path2MR-based histology localization with visually determined positioning using a hippocampus micro-atlas as reference^51^. The Path2MR coronal level of each histology section was obtained as the center of its gaussian distribution (width: σ=3mm), obtained around the predicted position to model registration error and registered to MNI. For the 72 Aβ histology sections included, resulting coronal MNI coordinates ranged between y=-8 (hippocampal head) and y=-42 (hippocampal tail). As shown in Figure 4A, these Path2MR coordinates presented a strongly significant correlation with visually determined anatomical positions (Pearson’s correlation, R=0.77, p=2.5x10^-15^). Additionally, we used the visual location of each section to generate a 3D position map using Path2MR, applying Equation 1 as explained in Figure 2 and Section 2.5:

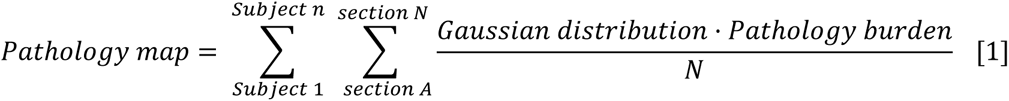

**Figure 4.**
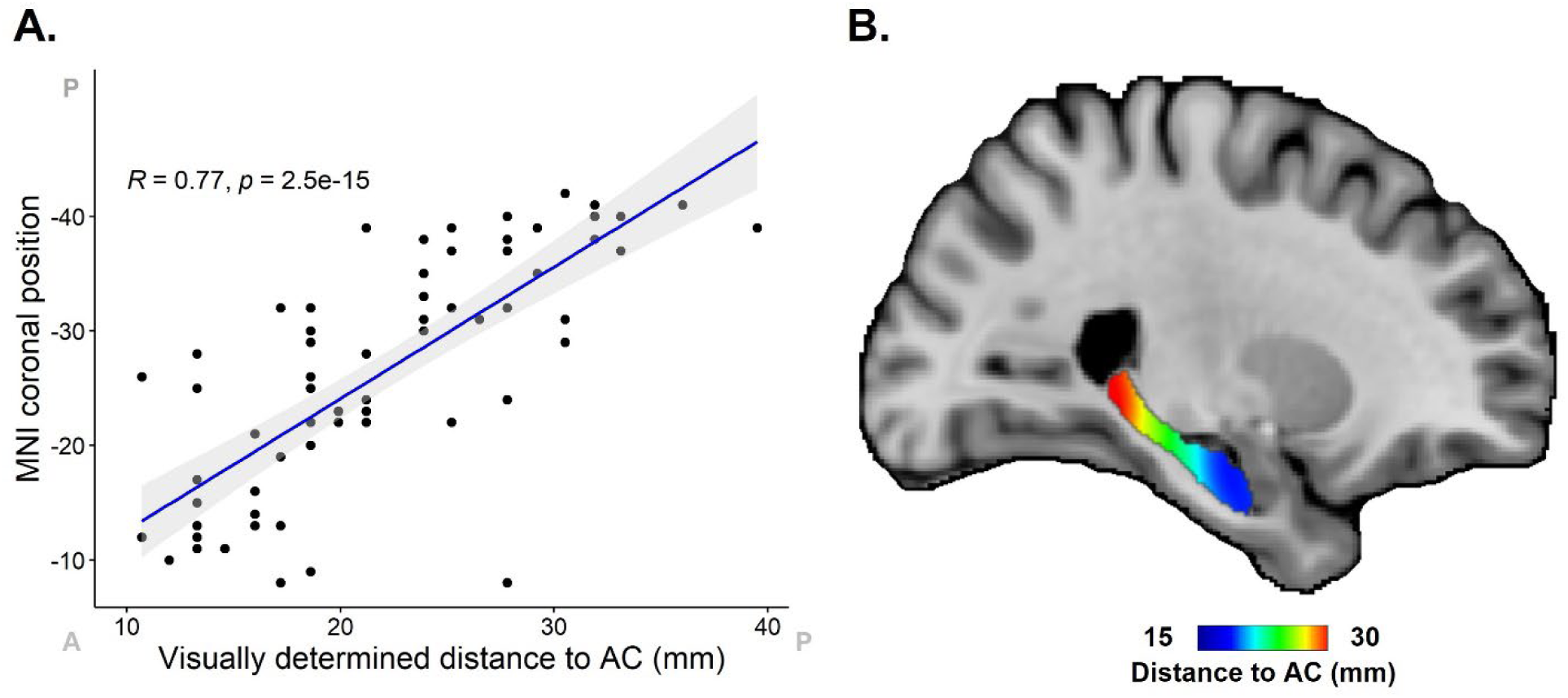
Comparison of histology localization with Path2MR with visually determined section positions. Visual positions were determined by using a micro-atlas of the hippocampus ^51^ as reference. **A.** Scatter plot and correlation between visually determined positions and coronal MNI position obtained from Path2MR. For each section, the slice with highest number of high-intensity voxels from the gaussian position distribution was used as its coronal MNI position. MNI coordinates have been reversed to ease visualization (from anterior to posterior). Anterior (A) and posterior (P) ends of x and y axes are indicated. Grey shaded region shows 95% confidence interval. **B.** Distribution map of distance to anterior commissure (AC) obtained with Path2MR, showing the shortest and largest distances in the hippocampal head and tail, respectively. AC: anterior commissure; MNI: Montreal Neurological Institute.

In this case, the visually determined distance to the AC was used as the “pathology burden” measure. The resulting hippocampal position map showed the expected distribution, with distance increasing progressively from the head to the tail of the hippocampus (Figure 4B). Both results were similar when using tau sections instead (R=0.77, p=5x10^-15^). Therefore, Path2MR conveys an anatomically consistent representation of histology localization, while overcoming the subjective character of visual comparison to a reference.

### 3.3 Anterior-posterior AD pathology distribution

The hippocampal long axis (anterior-posterior in humans) shows gradual and discrete transitions in terms of anatomical connectivity, genetics, receptor expression and pathology vulnerability^55^. To illustrate the utility of Path2MR for population analyses, we assessed the distribution along this axis of tau and Aβ pathologies, hallmarks of AD. For each pathology, we multiplied the gaussian distribution (σ=10mm) of each section by its quantified pathology burden, normalizing by the number of sections per subject and adding results from the 26 subjects (Equation 1, Section 2.5). Followed by applying a mask of the hippocampus, this strategy resulted in smooth pathology maps for tau and Aβ (Figure 5). For both pathologies, burden was highest at the hippocampal head, reaching lowest values at the hippocampal tail. To inspect these distributions closely, we obtained the mean intensity per coronal slice of each burden map, showing the highest tau burden at the anterior end of the hippocampus (MNI coordinate y=-7). Tau burden remained stable for the contiguous slices within the hippocampal head, followed by a linear decrease from y=-17 towards the posterior end of the hippocampus. In contrast, Aβ distribution resembled a quadratic curve, peaking at y=-18 within the hippocampal head. High-burden slices covered a more widespread region, with a decrease initiating at the hippocampal body and continuing throughout the tail.

**Figure 5.**
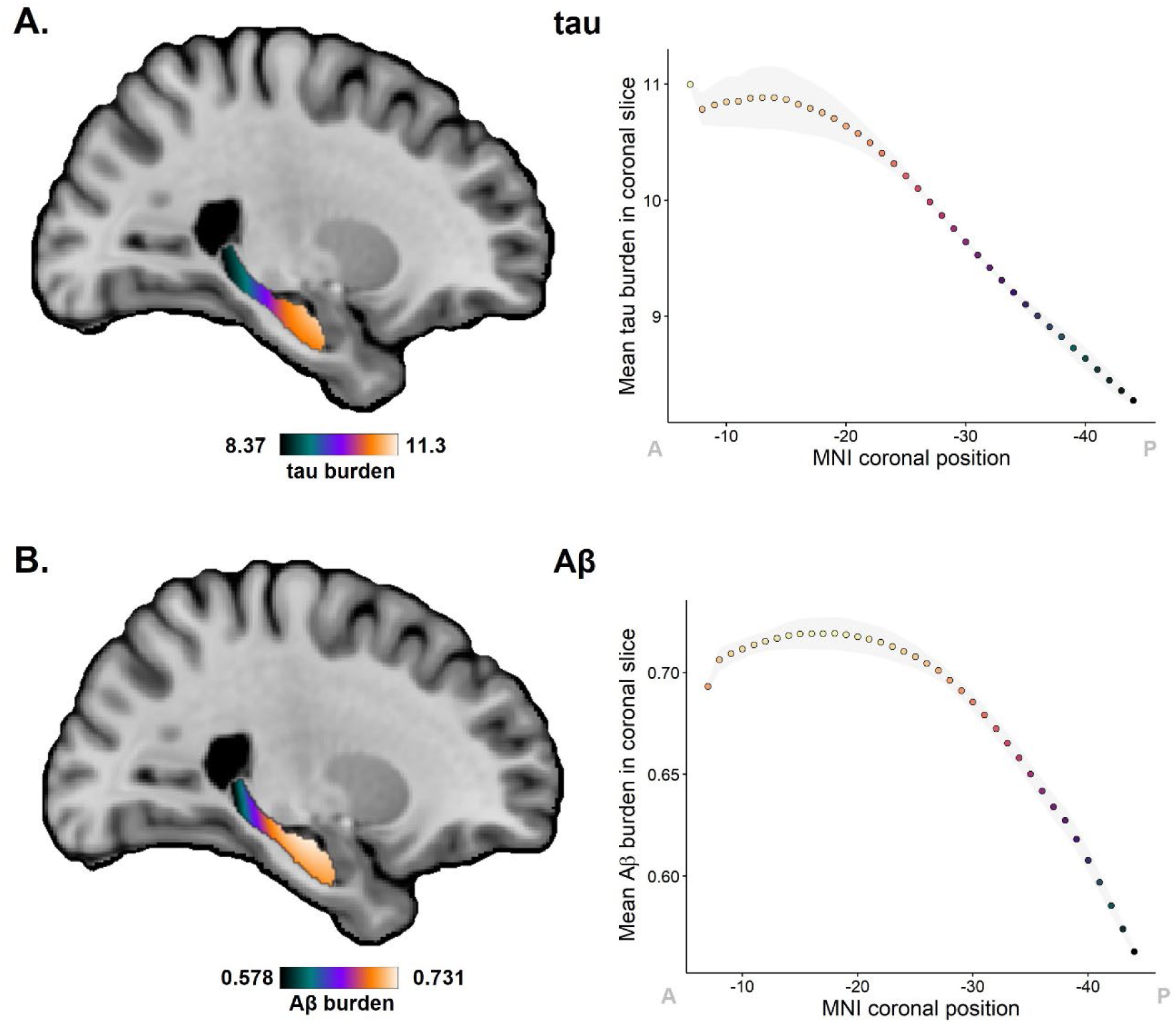
Distribution of tau and Aβ pathology burdens along the hippocampal long axis. **A.** Distribution map for tau pathology, quantified as the percentage of NFTs over the total neuron count, obtained with Path2MR. Mean tau burden per coronal slice of this map is shown in the right. **B.** Distribution map of Aβ burden, quantified as percentage area covered by Aβ plaques, together with plot of mean burden for every coronal slice. In graphs on the right, the grey shade shows standard deviation, and MNI coordinates have been reversed to ease visualization from anterior (A) to posterior (P). Aβ: amyloid-β; MNI: Montreal Neurological Institute.

These two anterior-posterior distributions were compared through a Kolmogorov-Smirnov test, showing a statistically significant difference between both (D=1, p=7x10^-16^). We also explored within-subject pathology gradients (Figure S2A), using MNI section positions obtained in Section 3.2. Given the low number of data points included per subject, individual pathology distributions were variable. However, in accordance with distributions from 3D pathology maps, a linear fit was found to be most appropriate for tau density, based on selection of fit with highest R^2^ and lowest AIC. Conversely, a quadratic function represented a better fit for Aβ pathology (Figure S2B). Results were similar when employing visually determined histology positions (Figure S2C), although this classification comprises a lower density of coronal levels through the hippocampus, and thus lower resolution. Therefore, Path2MR allowed population analyses at a finer resolution compared to a visual reference, revealing significant differences between tau and Aβ deposition along the long axis of the human hippocampus.

### 3.4 Pathology distributions for each subfield

Given that global section burden was obtained by averaging pathology values from five hippocampal subfields, we also evaluated differences across subfields. Mean tau (F(264)=248.4, p=7x10^-40^) and Aβ (F(295)=56.8, p=6x10^-13^) burden was significantly higher in CA1 and CA2 compared to CA3, LDG and MDG. Anterior-posterior differences were also found within each subfield for both pathologic proteins (Figure S3), as well as for separate measures of neuron and NFT areal densities (Figure S4). Neuron density in CA1, CA2 and CA3, and NFTs in all subfields, were higher at the anterior portion of the hippocampus, which explains the observed tau (NFT/neuron) distributions. We then generated subfield-specific 3D maps of tau and Aβ using Path2MR (Equation 1), from which the mean burden of every coronal slice was obtained (Figure 6). We found that, while all subfields presented their highest tau burden at the anterior end of the hippocampus, Aβ distribution was more variable across subfields. The anterior-posterior Aβ distribution of CA1, CA3 and LDG reached a maximum at the hippocampal head (MNI coordinates y=- 21, -7, and -8, respectively). In contrast, CA2 and MDG presented their maximum burden at the hippocampal body (y=-25 and -33, respectively). Therefore, the anterior-posterior gradient for Aβ pathology is intertwined with gradients in the proximo-distal axis (between subfields).

**Figure 6.**
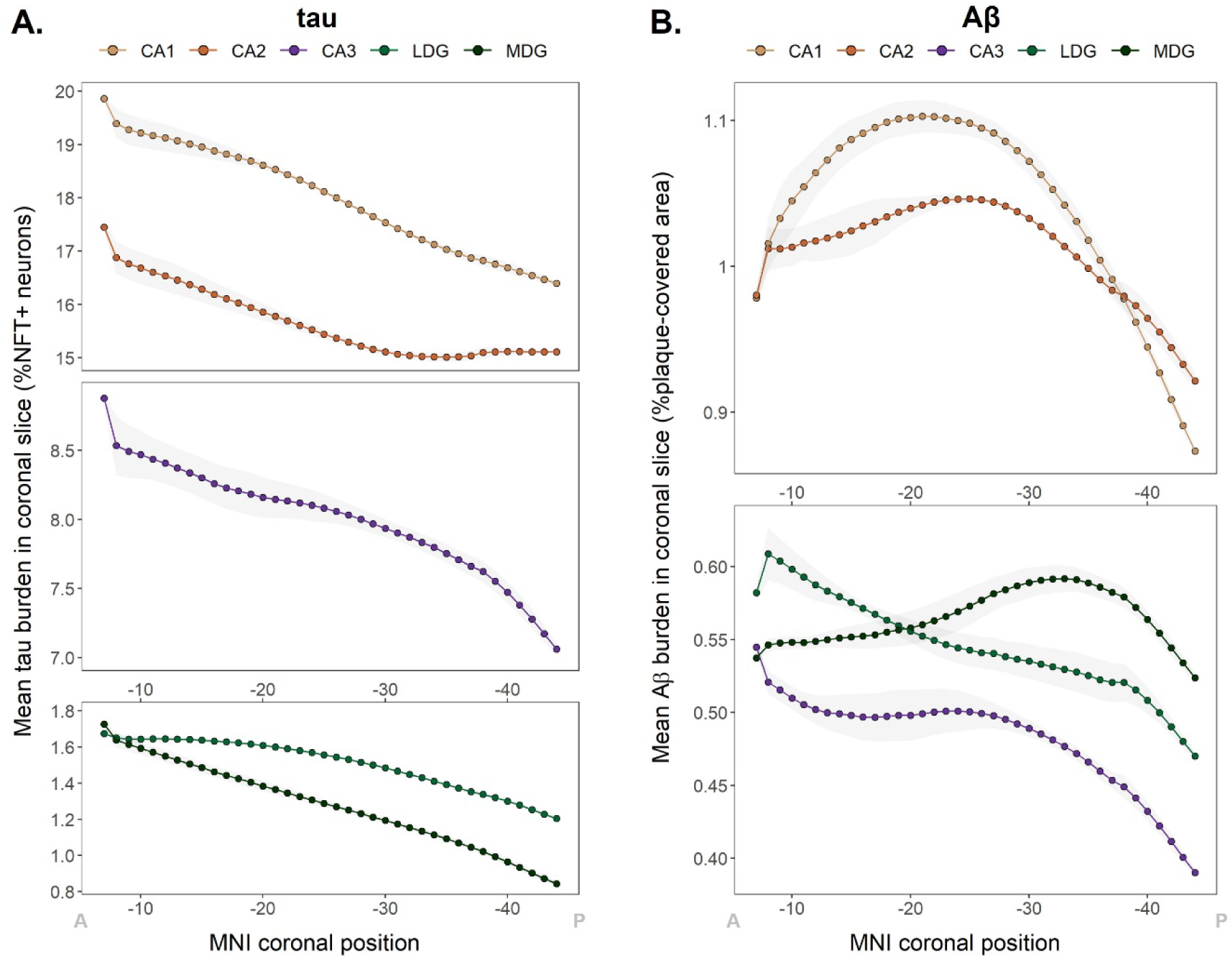
Individual distributions of tau and Aβ along the hippocampus for each subfield. **A.** Tau burden distribution obtained with Path2MR independently for each subfield, showing mean burden per coronal slice within the hippocampus. **B.** Mean Aβ burden in each subfield for every hippocampal coronal slice. Grey shades show standard deviation, and x axis coordinates are reversed for visualization from anterior (A) to posterior (P). Aβ: amyloid-β; LDG: lateral dentate gyrus; MDG: medial dentate gyrus; MNI: Montreal Neurological Institute; NFT: neurofibrillary tangles.

## 3 Discussion

We present Path2MR, a novel pipeline for histology mapping into a 3D framework enabling data integration for population analyses. This pipeline requires only blockface images of slices obtained upon sampling of the organ of interest – in this case, the left brain hemisphere. These images are stacked into a sparse 3D volume, which is then brought to isotropic 1mm^3^ resolution using deep learning and registered to atlas space using standard MRI processing methods. We implement Path2MR for population analysis using blockface images and hippocampal histology sections from 26 subjects with AD. In these experiments, the spatial interpretability of the method was verified by comparing histology mapping results in MNI space to visual section localization. While the MNI reference overcomes visual bias and provides a higher resolution compared to the anatomical micro-atlas (1 vs 1.4 mm), results from both methods showed a high correlation, and thus comparable outcomes. This supports the reliability of the method despite using specimens with severe atrophy. Moreover, Path2MR reconstruction, synthesis and MNI registration steps were quantitatively evaluated using ground truth data, revealing a 3D error of 3.27mm, which is within the previously reported 2-6mm error range from registration of brain MRIs to MNI^42,56,57^.

The critical advantage of our sparse histology reconstruction pipeline lies in its independence of a 3D reference image. Previous reconstruction methods were based on keystones obtained with highly specialized equipment and expertise: either a subject-specific MRI^26,27,58^, dense sampling of the specimen^9,12^, or both^21,23^. While dense sampling methods rely on stacking histological sections for reconstruction, Path2MR recovers 3D structure by stacking blockface images, which are straightforward to obtain. The proposed pipeline, however, entails several assumptions such as the depth of the histological section from the face of the tissue block. In contrast, direct stacking of serial histology sections allows considering different deformation sources during histology processing^59,60^. Using a specific 3D reference is also valuable to reduce uncertainty^8^, with sophisticated approaches such as mold-guided sectioning further facilitating the correspondence between MRI and histology^61,62^. Although Path2MR involves a larger registration error, we show it overcomes histological sampling variability for the hippocampus, which spans several centimeters along the coronal axis. As all steps of Path2MR are based on whole-brain methods, this approach can also be applied for histological analyses in any other brain structure, including cortical regions. This would be of great interest towards a more anatomically complete understanding of AD histopathological topographies.

The experiments included in this study revealed an anterior-posterior gradient of hippocampal tau NFT pathology. These results are in line with the recently reported higher anterior tau burden in the hippocampus and other MTL structures^31^, further supporting the reliability of the pipeline. In a PET study, tau radiotracer uptake was also found to be higher in anterior temporal regions^63^. The anatomical and functional connectivity between the human entorhinal cortex, where tau deposition initiates^3^, and the anterior hippocampus^55^, could provide insights into this increased regional burden. Interestingly, we also found a gradient of Aβ pathology which peaked towards the middle portion of the hippocampal long axis. To our knowledge, no other pathology studies have explored the distribution of Aβ along this axis. We also report intriguing neuronal density gradients in CA subfields of the hippocampus, with areal densities decreasing in the anterior-posterior direction. While these results are in contrast with previous studies in healthy subjects, which reported the opposite trend^64^, or no anterior-posterior differences^65^, new studies employing more recent methodology and including samples from AD patients are required to validate the gradients observed here.

Hippocampal anterior-posterior gradients have been consistently observed in other dimensions including gene expression^66^, as well as functional^55^ and anatomical^67^ connectivity. An additional axis of functional connectivity from the middle of the hippocampus towards anterior and posterior ends has been recently described using gradient mapping^68^. This gradient predicted episodic memory performance throughout the lifespan and could be linked to the Aβ distribution reported here. The topological distribution of amyloid PET binding has been shown to overlap with the “default mode” resting state network^69^. Evidence is emerging from analyses of functional gradients within the hippocampus and the cortical mantle that the default mode network may show strongest hippocampal connectivity at the transition between the head and body^70^. This has relevant implications for the understanding of clinical manifestations and progression of AD. In this context, it is pertinent to investigate hippocampal gradients of other proteopathies that commonly coexist with AD, such as TDP-43 and α-synuclein.

In line with previous studies^71,72^, we found both tau and Aβ burdens to be highest in hippocampal subfield CA1, followed by CA2. Differences between subfields were especially notable for Aβ, presenting a high variability in anterior-posterior distributions. This is coherent with the extracellular propagation pattern of Aβ, while tau spreads on a cell-to-cell basis through axonal connections^73^. As our analyses were performed by averaging burdens across subfields, whose relative positions differ along the long axis^51^, the obtained anterior-posterior distributions are conditioned by proximo-distal (between-subfield) differences. Recent approaches take advantage of histology digitalization for high-throughput quantification across the whole section, typically using deep learning^6,74^. Unfortunately, digitalization was not available for the studied dataset, limiting our ability to explore 3D topographical interactions in greater detail. In future work, Path2MR can be readily exploited using high-resolution digital histology data, to investigate pathological differences in medial-lateral and superior-inferior directions. This will provide a fine-grained understanding of spatial gradients within structures severely affected by neurodegenerative pathologies, as well as their associated cellular vulnerabilities.

Together with the lack of digital histology data, the presented experiments entail some limitations, including the unavailability of right-hemisphere pathological information. Another limitation is the use of blockface photographs acquired after histological processing. Brain slices used for reconstruction were non-intact, inducing uncertainty in subsequent steps of the pipeline. Kernel regression was therefore employed to model registration error, in a similar fashion to smoothing strategies typically used in neuroimaging analyses^75^. On the other hand, the robustness of the method to these experimental limitations is also a strength of this study, as it demonstrates the applicability of Path2MR to any brain bank dataset with access to preserved tissue. Its implementation with blockface photographs of intact brain slices will further improve the accuracy of its results, enabling more lenient error corrections. Another strength of this work is the use of spatial variability inherent to histological sampling to obtain a dense between-subject representation along the hippocampus, considering only 3 sections per subject. Combined with kernel regression, such dense representation over multiple subjects resulted in the smooth pathology maps presented here. Moreover, in our experiments, the possible effects of other co-pathologies on tau and Aβ patterns were isolated by studying a homogeneous set of patients with AD as the only substantial neuropathology.

In conclusion, Path2MR is a widely applicable 3D reconstruction pipeline. Using information extracted from routine dissection, Path2MR can be implemented with sparse histology and in the absence of an MRI reference. These features unlock the possibility to analyze in 3D the rich histological information obtained routinely at brain banks, as well as clinical and research institutions. We demonstrate the utility of Path2MR in population analyses including sections from three regions of the hippocampus (head, body and tail) from 26 AD patients. We found both tau NFTs and Aβ deposits predominate at the hippocampal head, while showing significantly different anterior-posterior distributions. Using this pipeline, future studies could validate these results and integrate them with data from earlier AD stages, as well as with other pathologies, to constitute a comprehensive map of combined hippocampal pathology in dementia.

## Consent Statement

All post-mortem donations included in this study were carried out under informed consent by a relative or proxy, following the BT-CIEN brain bank protocol approved by local health authorities.

## Acknowledgements

We greatly thank the donors of BT-CIEN and their families. We also thank the team of BT-CIEN, especially Paloma Ruiz for histology processing and staining. We thank Lidia Blazquez-Llorca and Yael Balbastre for helpful advice and discussions.

---

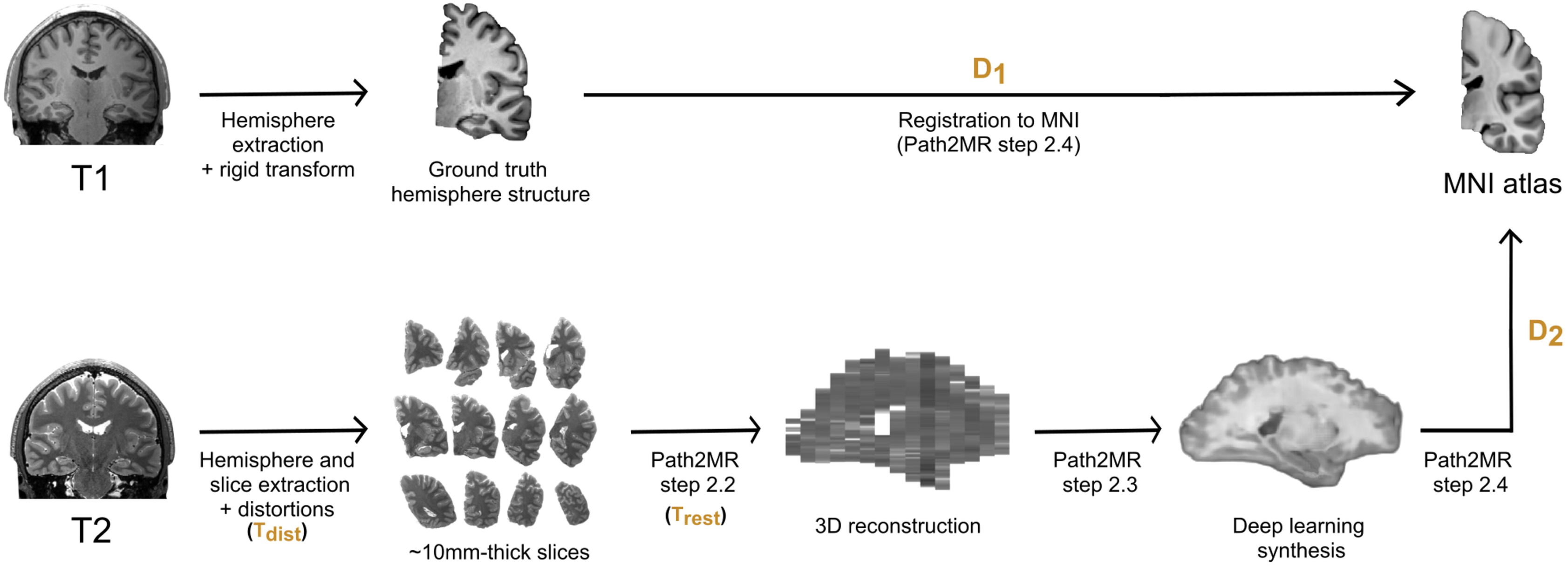

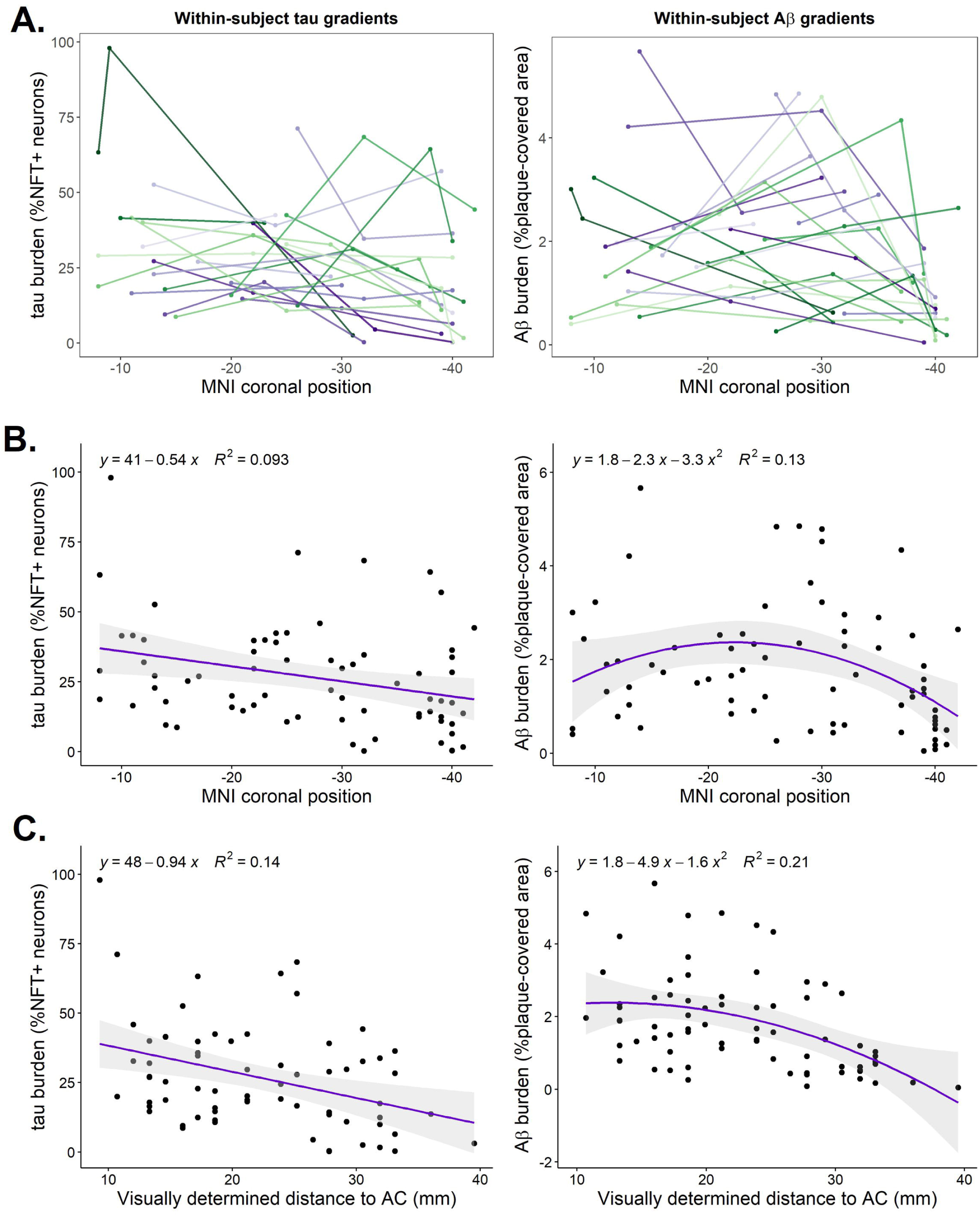

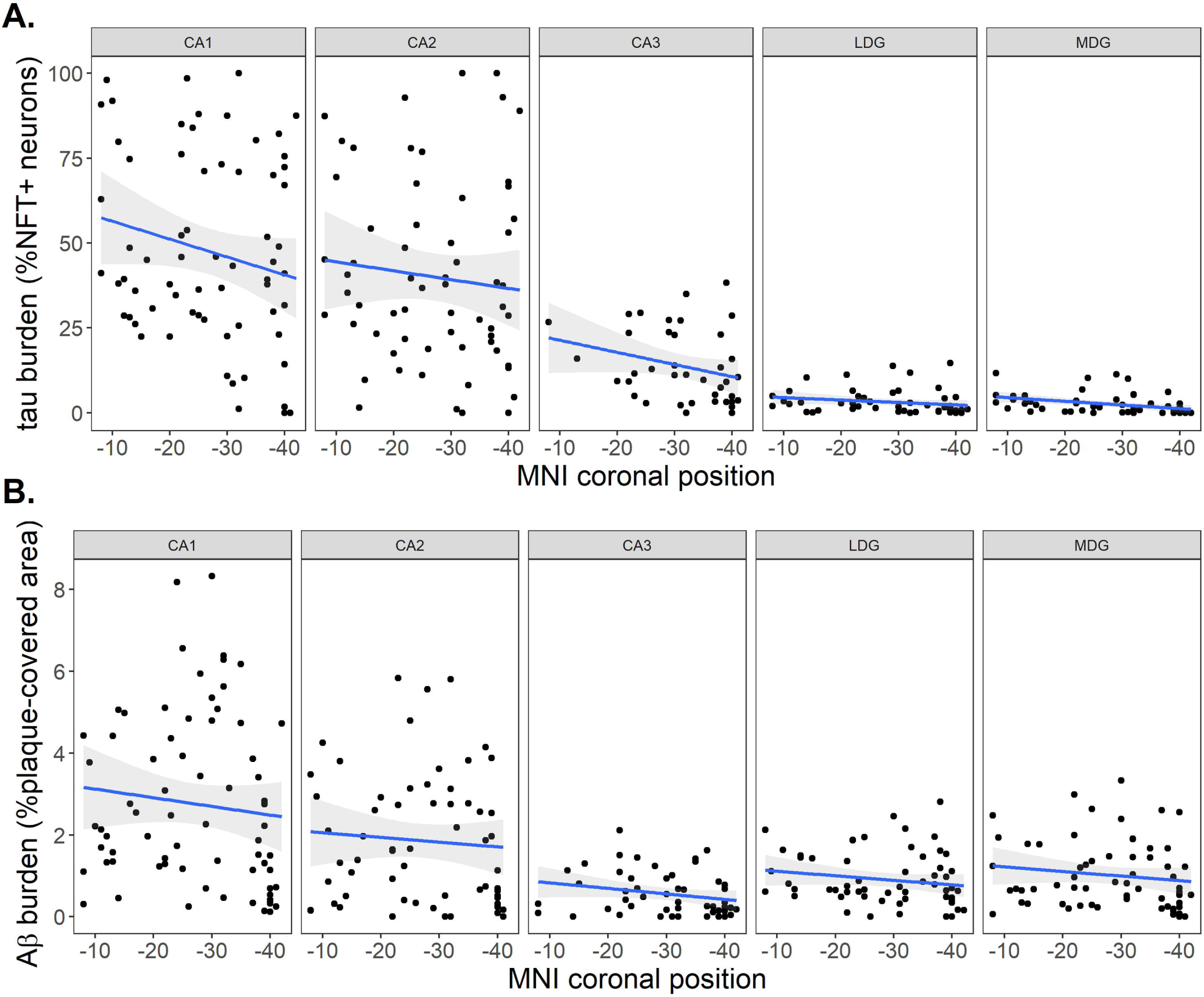

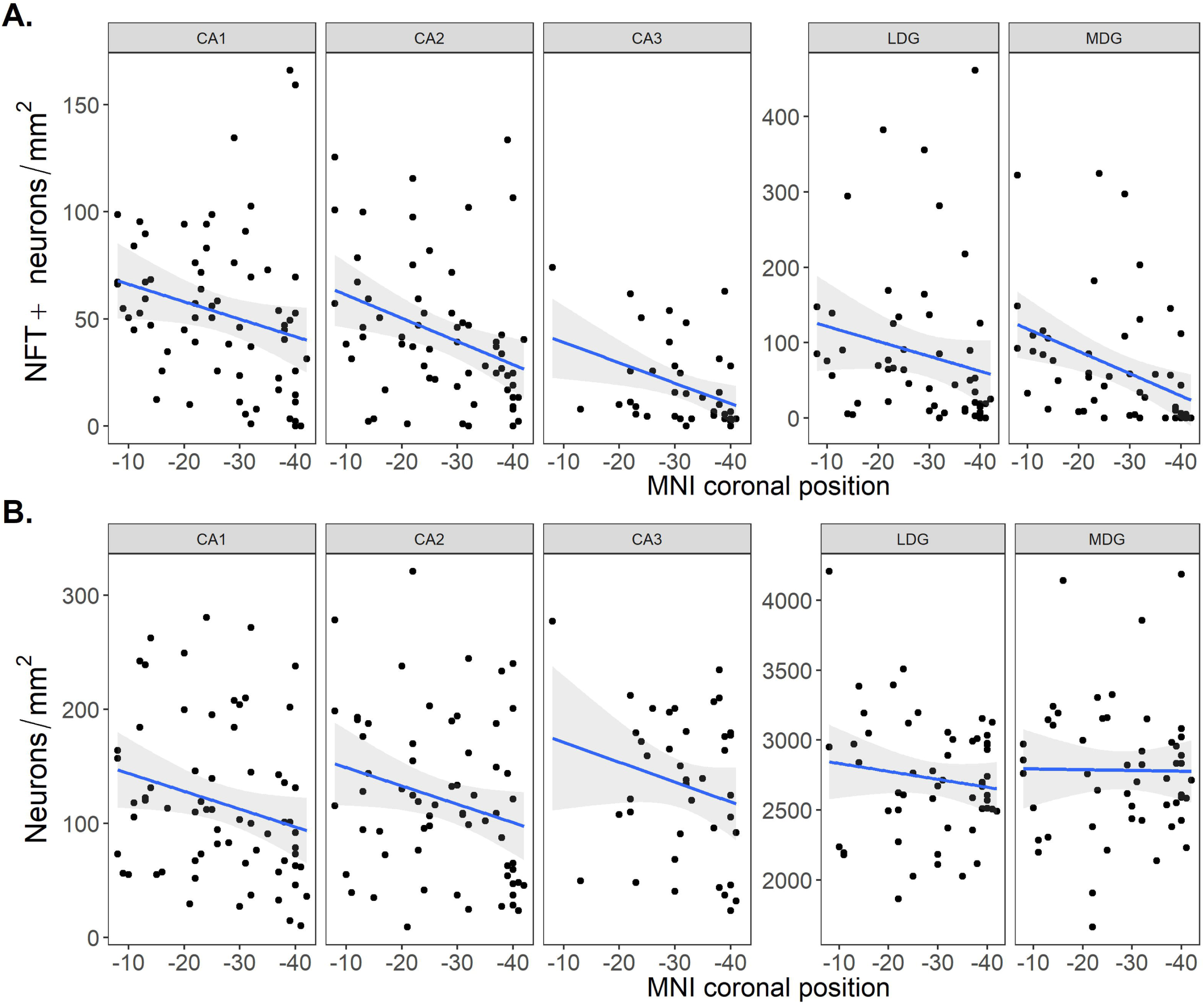

