## Supplementary Figures (S1-S4) for "Three-dimensional histology reveals dissociable human hippocampal long axis gradients of Alzheimer’s pathology"

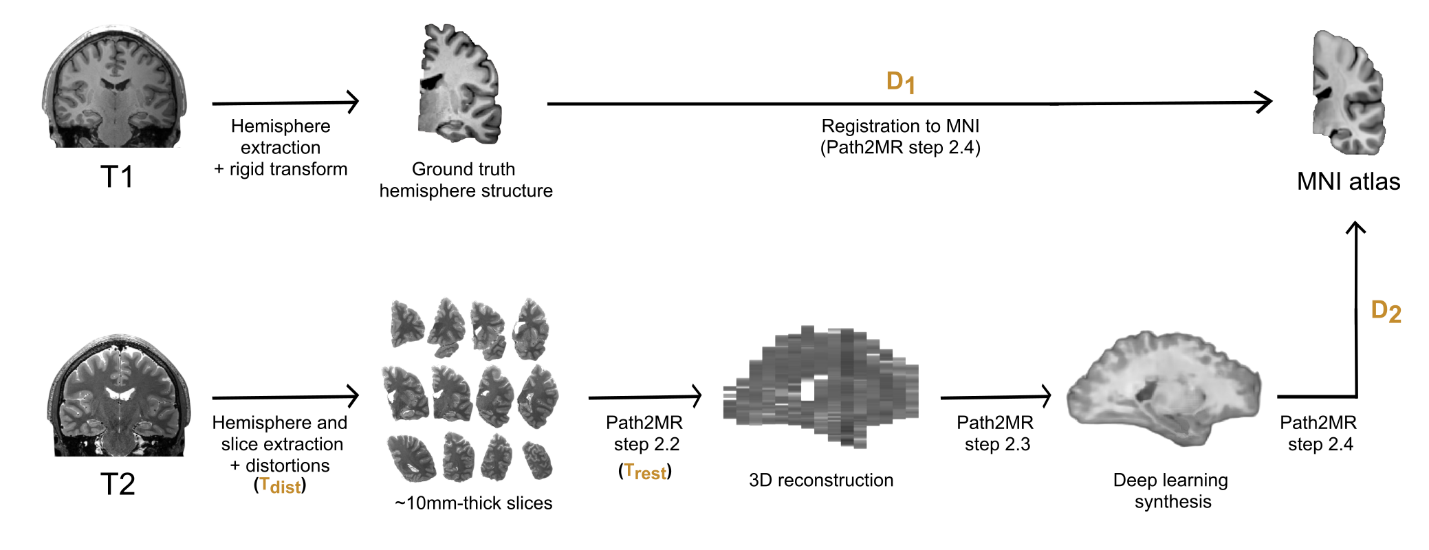

**Figure S1. Flowchart of validation experiment of Path2MR reconstruction performance using the HCP dataset.** T1 and T2 scans from 100 subjects were retrieved, with the former being used as ground truth reference, and the latter being used for slice simulation and Path2MR processing. Random transformations were applied to the T1 (3D rigid transforms) and T2-derived slices (2D distortions, T_dist_). After Path2MR reconstruction (restoration transform T_rest_) and synthesis, both the outcome and the T1 ground truth were registered to MNI, and the error was obtained as the difference between resulting deformations (D_1_ and D_2_, respectively). 3D: three-dimensional; HCP: Human Connectome Project; MNI: Montreal Neurological Institute.

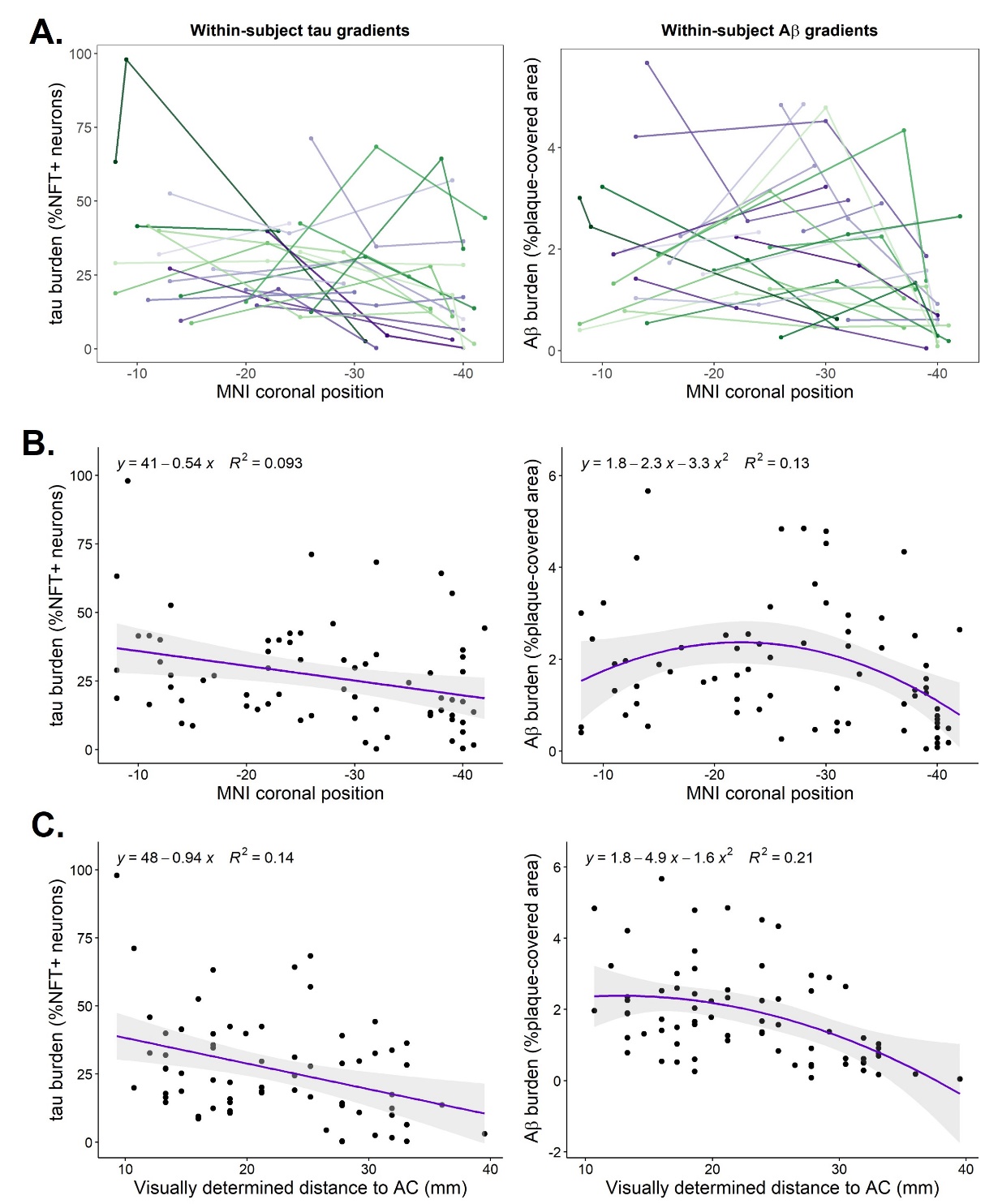

**Figure S2. Distribution of tau and Aβ pathologies along the hippocampal long axis. A.** Within-subject pathology gradients for tau (left) and Aβ (right), with lines connecting hippocampal sections from the same subject. For tau analyses, two subjects had only one section included and were not considered in this plot (n=24). **B.** Data points from tau and Aβ sections from all subjects as a function of MNI positions derived from Path2MR. Linear fit is shown for tau, which significantly explained these data (p=.011), while Aβ distribution was better predicted by a quadratic fit (p=.008). MNI coronal coordinates in the x axis have been reversed to ease visualization (from anterior to posterior). **C.** Tau and Aβ burdens from all subjects as a function of visually determined section positions, again showing a better fit with a linear equation for tau (p=.002) and quadratic for Aβ (p=3·10^-4^). Aβ: amyloid-β; AC: anterior commissure; MNI: Montreal Neurological Institute; NFT: neurofibrillary tangles.

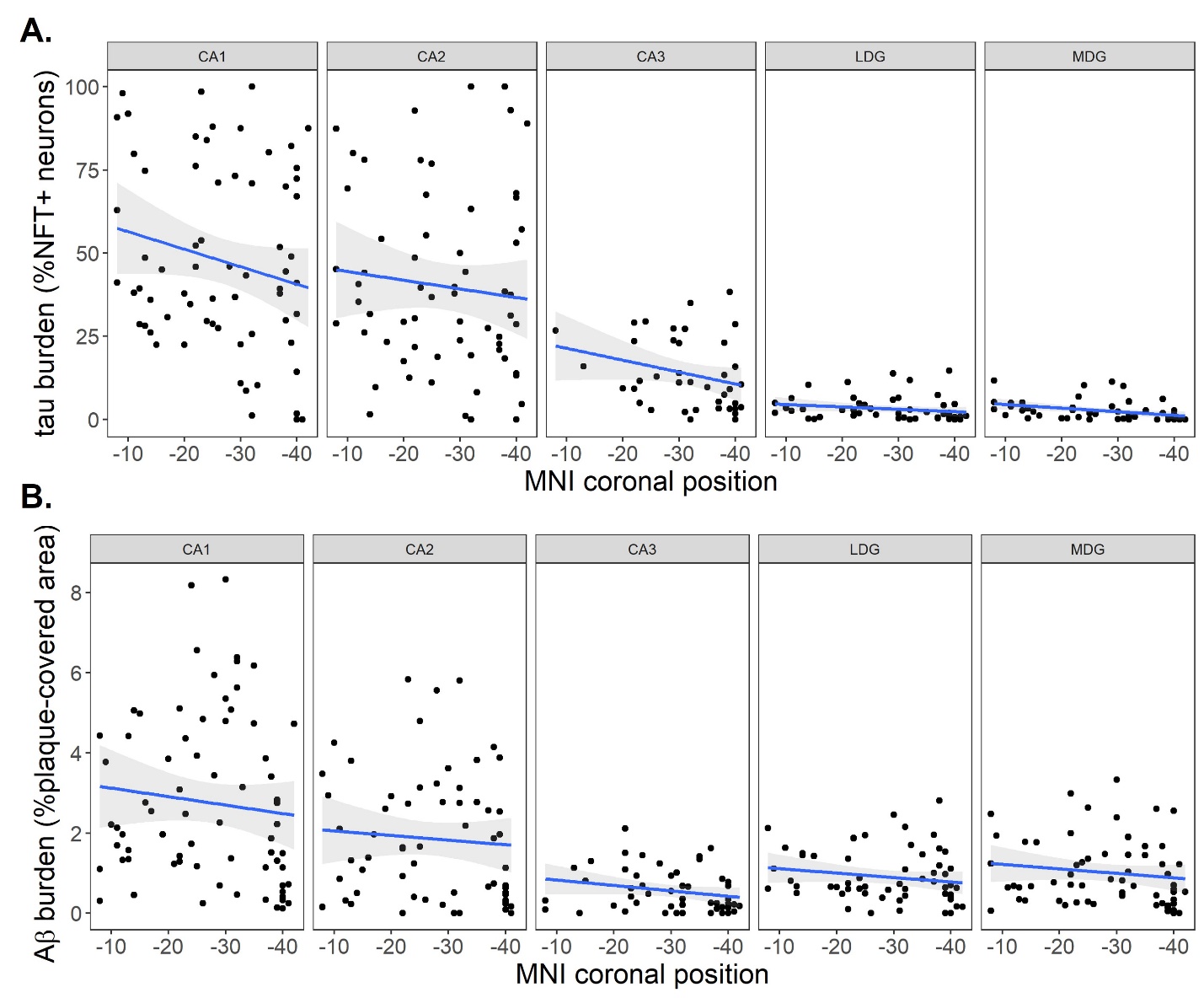

**Figure S3. Subfield differences in tau and Aβ deposits along the hippocampal long axis. A.** Separate tau quantification (%NFT+ neurons) results for each subfield, as a function of MNI positions derived from Path2MR. **B.** Quantification results of Aβ images for each subfield, as a function of MNI positions from Path2MR. Coordinates of the x axis have been reversed for visualization from anterior to posterior. Both results are similar when using visually determined section positions instead of those from Path2MR. Linear fits are added to ease visualization. Aβ: amyloid-β; MNI: Montreal Neurological Institute; NFT: neurofibrillary tangles.

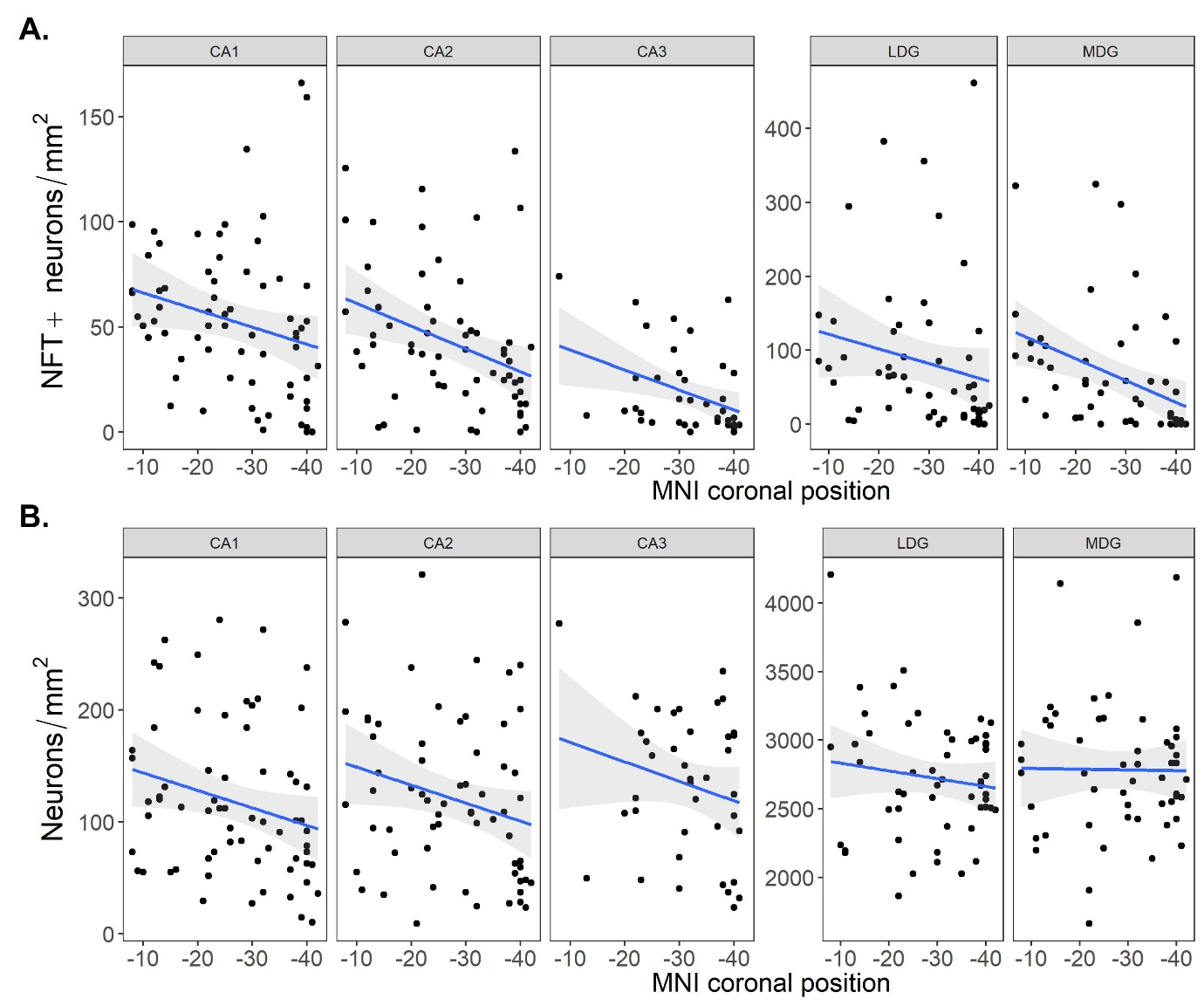
 **Figure S4. Differences in separate measures obtained for tau quantification, between subfields and along the hippocampal long axis. A.** NFT+ neuron count per mm^2^ for each subfield as a function of the position of each histology section derived from Path2MR (N=276). **B.** Total neuron (including NFT- and NFT+ ones) areal density within each subfield as a function of section positions from Path2MR (N=276). Both (A) and (B) are similar when using visually determined section positions instead of those from Path2MR. Coordinates in the x axis have been reversed for visualization from anterior to posterior. In CA fields, neuron counts are normalized by pyramidal layer area (covering the whole image), while in DG fields they are normalized by granular layer area (measured separately). Linear fits are added to ease visualization. MNI: Montreal Neurological Institute; NFT: neurofibrillary tangles.
